# “Value” emerges from imperfect memory

**DOI:** 10.1101/2024.05.26.595970

**Authors:** Jorge Ramírez-Ruiz, R. Becket Ebitz

## Abstract

Whereas computational models of value-based decision-making generally assume that past rewards are perfectly remembered, biological brains regularly forget, fail to encode, or misremember past events. Here, we ask how realistic memory retrieval errors would affect decision-making. We build a simple decision-making model that systematically misremembers the timing of past rewards but performs no other value computations. We call these agents “Imperfect Memory Programs” (IMPs) and their single free parameter optimizes the trade-off between the magnitude of error and the complexity of imperfect recall. Surprisingly, we found that IMPs perform better than a simple agent with perfect memory in multiple classic decision-making tasks. IMPs also generated multiple behavioral signatures of value-based decision-making without ever calculating value. These results suggest that mnemonic errors (1) can improve, rather than impair decision-making, and (2) provide a plausible alternative explanation for some behavioral correlates of “value”.

## 1 Introduction

The brain is an imperfect computer. There are fundamental constraints on its ability to accurately represent, retain, and recall the information we have encountered in the world [1–3]. While neural processing constraints can cause errors that have no obvious utility [4], others may generate “errors” that are computationally useful [2, 5, 6]. Especially in uncertain environments, there is both theoretical [7] and empirical [8, 5] evidence that noise is critical for exploratory discovery and learning. By contrast, in cognitive models, noise is generally only added as an error term, with limited but important exceptions [9, 10, 2]. Developing cognitive models that meaningfully incorporate noise could be the key to determining when the constraints on the brain are truly a limitation and when they serve a computational purpose.

This paper offers a thought experiment: What if neural processing errors were sufficient for making good decisions in uncertain environments? Sequential decision-making in changing and only partially observable environments requires continual learning and discovery. This is exactly the kind of task where computational noise is most likely to be useful [8, 9]. Yet realistic errors in representing, retaining, or recalling past rewards are not a feature of cognitive models of sequential decision-making. Instead, the standard approach assumes that previous rewards are perfectly encoded and integrated into a value signal that is then used to guide decisions [11–13]. While some recent work has considered alternative architectures, including sampling from episodic memory [14–16], these studies have not permitted memory errors. As a result, we know little about how errors would impact decision making, nor how an agent imbued with this noise would differ from other models or real decision-makers.

To address this omission, we developed a novel and minimal decision-making agent that we call “Imperfect Memory Programs” (IMPs). IMPs systematically misremember the timing of past rewards in a biologically realistic way, but perform none of the value computations thought to be important for sequential decision-making in the cognitive sciences. We find that IMPs are capable of performing well above chance in multiple decision-making environments and generate behavior that is typically interpreted as evidence of value integration. These results have implications for understanding sequential-decision making, but also for designing new algorithms for decision-making in uncertain environments.

## 2 Imperfect Memory Programs (IMPs)

IMPs (Fig. 1A) use a 2-step process to make decisions. First, in the sampling stage, IMPs “remember” a past outcome via a noisy sampling process. Second, in the choice stage, IMPs decide whether to exploit the previous action or to explore the action space.

**Fig. 1.**
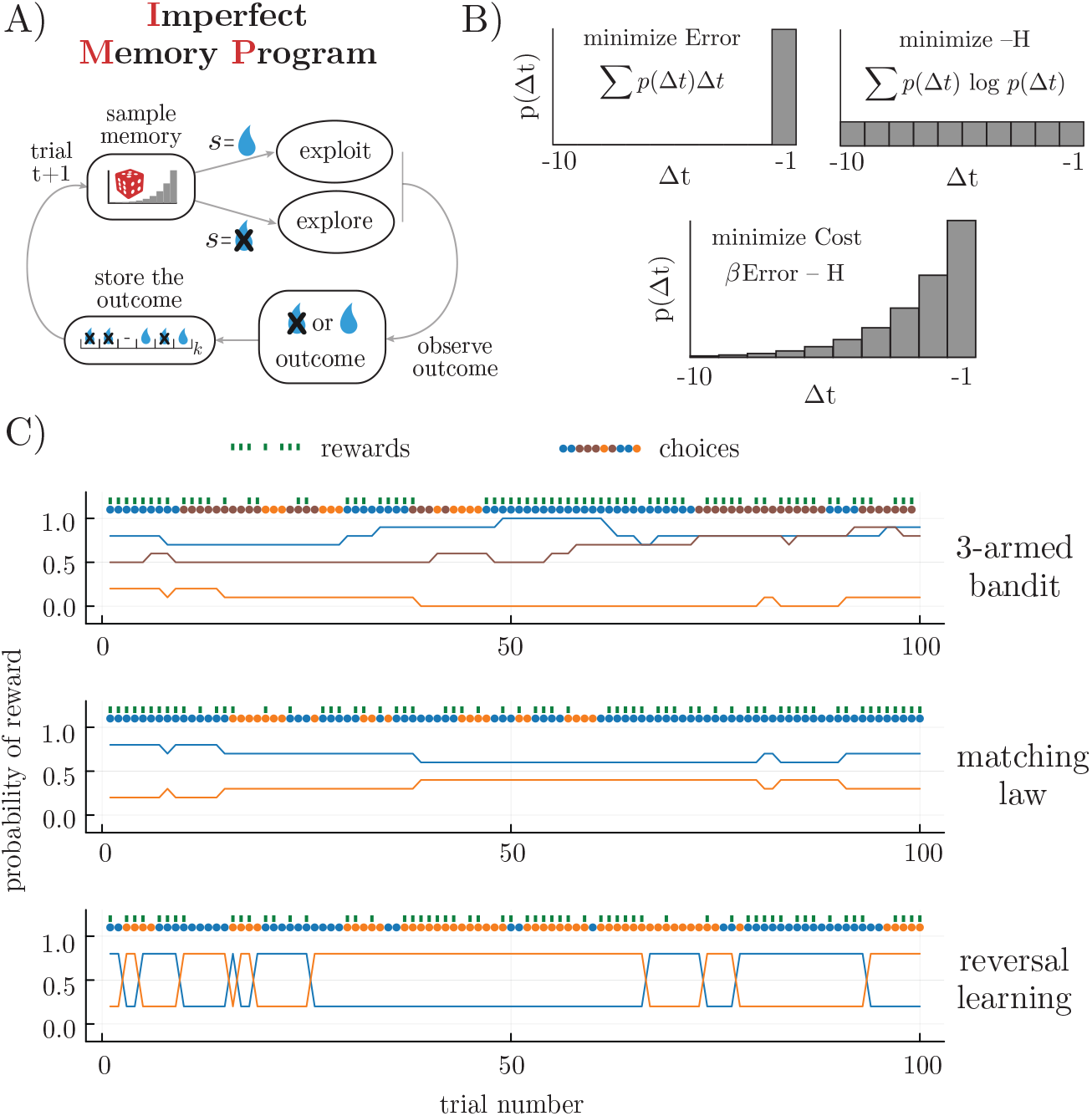
Agents and environments. A) Schematic illustrating the structure of the decision-making process in Imperfect Memory Programs (IMPs). B) IMPs sample previous rewards from memory in a way that optimizes the trade-off between (1) minimizing the average temporal error of the recalled samples (top left) and (2) minimizing the complexity of the sampling process (top right). The Boltzman distribution (bottom) minimizes the total cost (or free energy) of the sampling distribution, thereby naturally balancing these two objectives. Here *β* is both the inverse temperature of the sampling process and the single free parameter of the IMPs. C) Examples of the reward schedules from 3 testbeds: a restless 3-armed bandit (top), a matching law task (middle), and a probabilistic reversal learning task (bottom). Delivered rewards and the choices generated by one example IMP are overlaid.

In the **sampling stage**, IMPs sample a reward outcome from a memory store that contains the set of previous rewards associated with the current action. That is, they remember some prior reward *R* from a previous time point *Δt ∈* (*−∞*, 0]. Inspired by previous theoretical work in the domain of structure learning [2], we assume that this memory store is organized such that it balances 2 competing objectives (Fig. 1B). First, the memory store should maximize the likelihood of recalling relevant past outcomes. In changing environments, the relevance of past information decreases as a function of time, such that recalling outcomes distant in time (*Δt* with respect to the present), incurs a high error. The sampling distribution *Q* that would maximize timeliness is the one that minimizes the average temporal error in remembering, *E*(*Q*) = ∑_*Δt*_ *Q*(*Δt*)*Δt*. Second, the recall process should be as minimally complex as possible (i.e., efficient to implement), while also maximizing the information that can be sampled from memory. This fact implies that we should choose the maximum entropy sampling distribution *Q*: the one that minimizes the negative entropy, *−H*(*Q*) = ∑_*Δt*_ *Q*(*Δt*) log *Q*(*Δt*). We can balance these two competing objectives by calculating the total cost, or free energy of *Q, F* (*Q*) = *βE*(*Q*) *− H*(*Q*), and minimizing *F* (*Q*) with respect to *Q*. In doing so, we find that the sampling distribution that would simultaneously minimize the magnitude of mnemonic errors and maximize information storage is the Boltzman distribution,

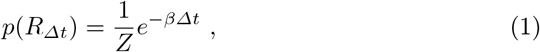

where *Z* is the partition function and *β* is the model’s single free parameter, which controls the trade-off between error magnitude and storage capacity. Although this choice is theoretically motivated [2, 17], there is also empirical evidence that memory retrieval tends to be exponentially recency-weighted [1, 2, 18], suggesting that the brain may also have struck a balance between error magnitude and complexity. In instances where IMPs are unable to sample a past outcome from memory (for example, during the first choice to an option), an outcome (reward omission or delivery) is chosen at random as the sample.

In the **choice stage**, IMPs make a deterministic choice based on the sampled outcome. Inspired by an *ϵ*-greedy decision rule, IMPs choose whether to explore or exploit. If the remembered outcome, *R*_*Δt*_ was not positive (not rewarded), the agent explores through choosing a new action policy at random,

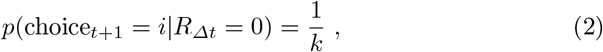

where *k* is the total number of options. If the remembered outcome was positive (rewarded), the agent continues to exploit its current action policy,

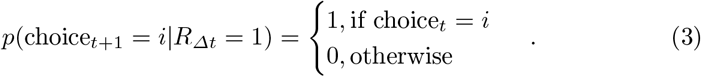

This decision policy was chosen for its simplicity, but there is also evidence that biological decision-makers produce distinct explore and exploit choices in similar tasks, where the exploration resembles random decision-making and exploitation resembles directed, reward-dependent decision-making [19–21].

## 3 Testbeds

We simulated behavior from the IMP in 3 sequential value-based decision-making tasks that are common in the neuroscience and psychology literature (Fig. 1C). These included a restless bandit task [19, 22, 23, 20, 5], a matching law task [24], and a probabilistic reversal learning task [25, 26]. Unless otherwise noted, simulations involved 500 sessions of 500 trials. All agents experienced identical environments.

In each task, choices are made between a set of *k* options, each of which is associated with some probability of reward. Reward probabilities can only be inferred by choosing each option and combining information over multiple samples. The tasks are all uncertain because the reward probabilities are not fixed, but instead evolve over time. This encourages decision-makers to exploit valuables option when they are discovered while also occasionally exploring alternative options that have the potential to become more rewarding at any time.

In the **restless bandit task** (Fig. 1C, top), the reward probabilities of each option *i* are independently updated at each trial *t* according to

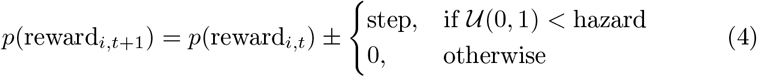

where “hazard” is a fixed rate of change ∈ [0, 1], 𝒰 (0, 1) is a draw from a uniform random distribution, and the sign of the step is chosen independently at random for each option on each trial. The hazard rate and step size were both fixed at 0.1, and the number of options was fixed at 3 except as otherwise noted, after [21, 5, 6, 26].

In the **matching law task** (Fig. 1C, middle), reward probabilities are updated according to the same function, but not independently because

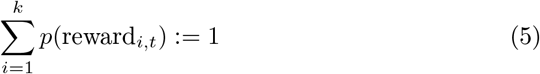

for each option *i*. The matching law task is often used to illustrate that biological decision-makers tend to be imperfect reward maximizers: more likely to allocate their choices in proportion to the rate of reward than to chose the best option. Matching law tasks are typically 2-alternative; we followed that convention here. The **probabilistic reversal learning task** (Fig. 1C, bottom) is another 2-alternative decision-making task that is very common in rodents. As in the matching law task, reward probabilities are symmetrical such that one option is high-value and the other is low-value. However, here the high and low values are fixed, often at *p*(reward|high) = 80% and *p*(reward|low) 20%, with the identity of the high and low values swapping at specific reversal points.

## 4 Results

### 4.1 IMPs outperformed simple agents with perfect memory

Simulations in a restless k-armed bandit environment revealed how IMPs behavior and performance depends on the inverse temperature parameter *β*. When *β ≫* 1, the cost of recall error takes over the total error, and IMP agents become perfect memory agents with a Win-Stay, Lose-Shift (WSLS) strategy (Fig. 2A-B, right). In contrast, when *β ≪* 1, the complexity cost dominates and IMPs stochastically recall reward episodes far in the past; hence, performance degrades in this unpredictable environment (Fig. 2A-B, left). Crucially, there is an intermediate value of *β* that not only optimizes the performance of the IMPs and outperforms WSLS (Fig. 2A-B), but does so while switching less frequently (Fig. 2C). Given that switch decisions take longer than stay decisions [27], IMP’s tendency to repeat choices suggests that a biological agent would be able to achieve a higher rate of reward via an IMP-like algorithm than a WSLS-algorithm.

**Fig. 2.**
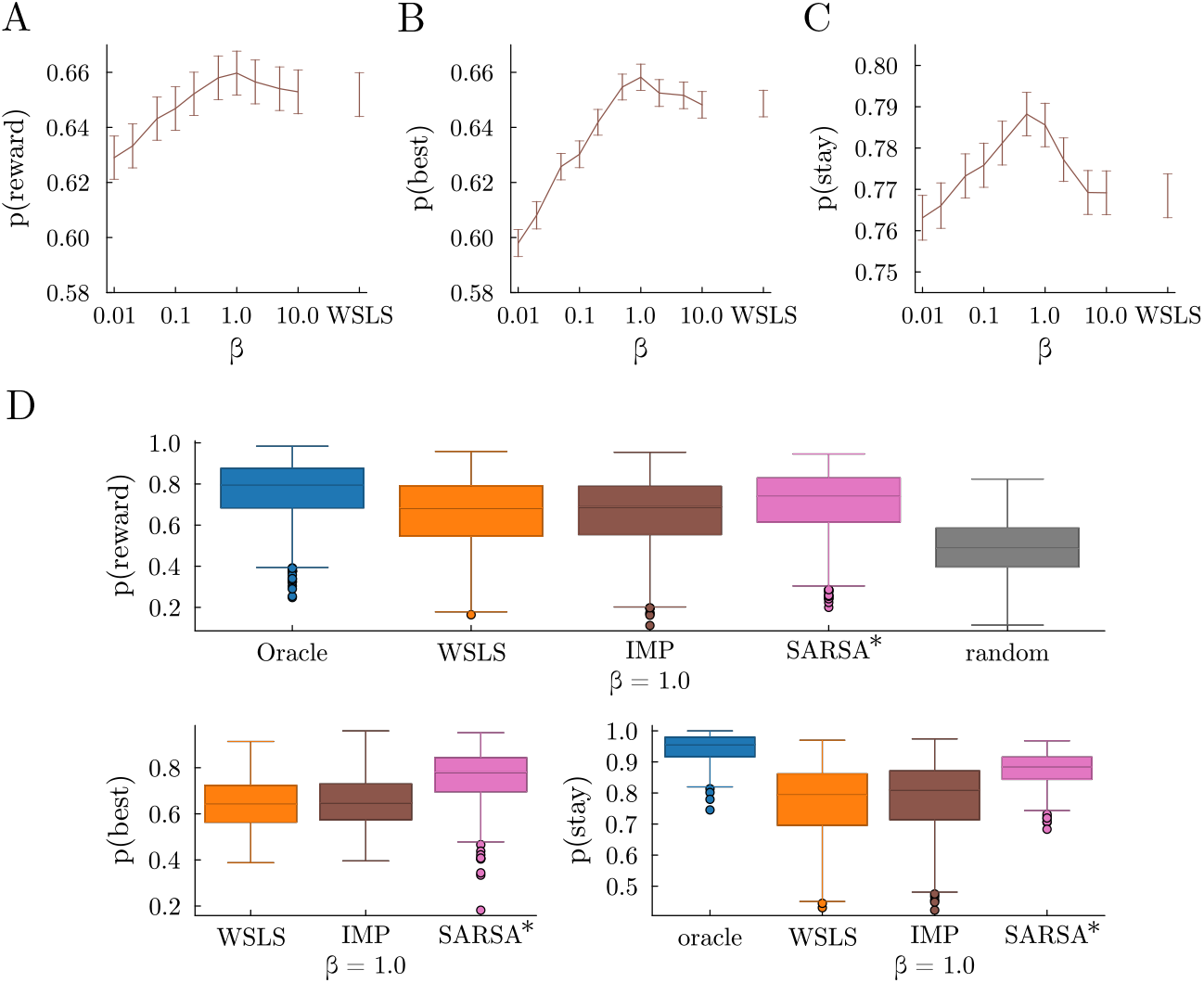
IMPs perform a restless multi-armed bandit task without perfect memory. Probability of (A) obtaining reward, (B) choosing the best arm, and (C) sticking with the same arm across sessions, as a function of the inverse temperature *β* that trades off complexity and error. WSLS is a special case of an IMP with perfect memory, corresponding to infinite *β*. (D) Probability of obtaining reward, choosing the best arm (or any of the best arms) and of repeating a choice for various agents, with their free parameters optimized for this task.

To benchmark IMPs’ performance, we also simulated an oracle (that knows the probability of each arm and always selects the best), a random agent (which chooses an arm uniformly randomly), and a reinforcement learning algorithm (SARSA, [11]). SARSA updates the value *Q*(*a*) of an arm *a* after receiving reward *R* at time *t*, and sampling another arm *a*^*′*^ from its policy,

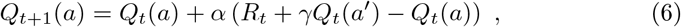

where *α* is a learning rate and *γ* is a discount factor. Then, the SARSA agent defines a probability *π*(*a*) of choosing an arm *a* using the action value *Q*(*a*), as a softmax distribution

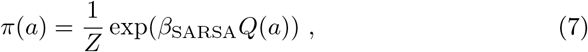

where *β*_SARSA_ is an inverse temperature parameter that controls the noise for the SARSA agent, and *Z* is a normalizing function.

While the optimal IMPs did slightly better than WSLS agents in the restless arm bandit task, they still underperformed SARSA, with optimized parameters *α* = 0.8, *γ* = 0.9, and fixed *β*_SARSA_ = 10 (Fig. 2D), and all of these agents perform better than chance. For the parameters of this task, both the oracle and the optimized SARSA agent show that staying persistently is a good strategy to obtain more rewards (Fig. 2D, probability of staying is higher for these agents), which the IMPs tend to do more than perfect memory WSLS agents.

### 4.2 IMPs generate reward history kernels

Sequential decision-making algorithms like SARSA, generally assume that agents calculate a quantity known as “value” by integrating experienced rewards. By setting the discount factor *γ* to 0 and rearranging Eqn. 6, we see that SARSA is one example of the broad class of delta-rule learning models where

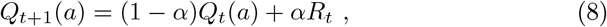

which makes explicit that action values *Q*(*a*) are the *α*-weighted average of value at the previous time step and the newest reward. This implies that the weight of past rewards falls off exponentially in these models, mirroring the pattern that is commonly seen in biological decision-makers [28].

IMPs do not calculate value. Nonetheless, it seems plausible that they might generate value-like reward history kernels because of their memory swaps. To measure their reward history kernels, we simulated IMPs in a 2-armed bandit and fit a logistic regression model,

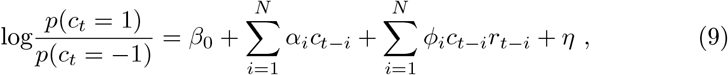

where *c*_*t−i*_ is 1 if the first option is chosen on trial *t−i* (−1 if the second is chosen), and *r*_*t−i*_ is 0 if they were rewarded on that trial (0 otherwise). Together, the *ϕ*_1:*N*_ parameters represent the unique effect of previous rewards on the log odds of choice, beyond the contribution of choice history (*α*_1:*N*_) and bias (*β*_0_). Models were fit via ridge-regularized maximum likelihood (*λ* = 1). To determine if the influence of previous rewards decayed exponentially quickly, we fit a 3-parameter exponential curve, *Ae*^*−Bx*^ *− C*, to *ϕ*_1:*N*_. Here, *A* represents a scaling parameter, *B* is the decay rate of the influence of previous rewards, and *C* is an offset.

IMPs reliably generated reward history kernels that were well-described by exponential decay (Fig. 3A-C; median *R*^2^ = 0.9999). The decay in the reward history kernel largely increased as a function of *β* before saturating at values *>* 5 (Fig. 3D). In short, IMPs had exponentially decaying reward history kernels because of their imperfect memory, rather than any value calculations.

**Fig. 3.**
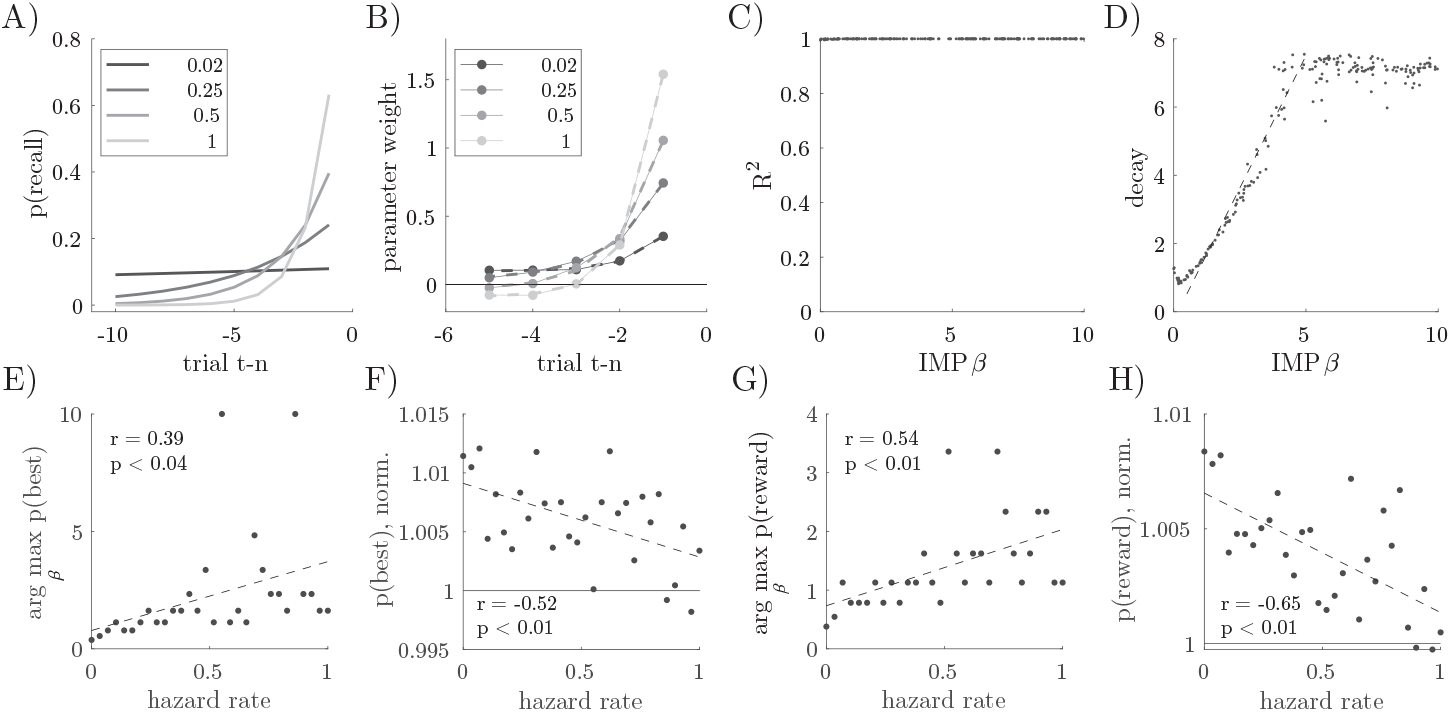
IMPs integrate reward history via imperfect memory. A) Probability of recall as a function of previous trial for 4 example IMPs with different *β*. B) Reward history kernels for the 4 example IMPs, with exponential fits overlaid. C) *R*^2^ as a function of *β* for 200 simulated IMPs with *β ∼ U* (10^*−*2^, 10^1^). D) Exponential decay parameter for the simulated IMPs in panel C. Gray line is the least squares linear fit for 0.5 *≥ β ≤* 5 (slope = 1.54, offset = -0.23). E) The *β* that maximizes the probability of choosing the best option, plotted as a function of volatility (hazard), identified via grid search (20 log-distributed bins ∈ [10^*−*2^, 10^1^]). F) Probability of choosing the best option for the optimal IMP, plotted as a function of volatility. *P* (best) is normalized to WSLS performance at each volatility level (WSLS at 1). G-H) Same as panels E-F, for *p*(reward).

If IMPs imperfect memory accomplishes something like reward integration, then in less volatile environments, where longer reward history integration offers an advantage, (1) IMPs should have a greater advantage over WSLS, and (2) the optimal *β* should decrease. To test these predictions, we simulated IMPs in the restless bandit with varying hazard rates. We found that the optimal *β* and the performance of the optimal agent both scaled with volatility (Fig. 3E-H). This observation suggests that the imperfect memory process in IMPs functioned like the reward history integration in delta-rule learning agents.

### 4.3 IMPs engage in behaviors between matching and maximizing

Under some circumstances, biological decision-makers tend to match the relative rate of reward of their options rather than maximize their reward by consistently choosing the best option [24, 29–32]. In order to determine whether IMPs tended to match or maximize, we simulated IMPs in a matching law testbed. We found that IMPs tended to match: allocating their choices in proportion to the rate of reward associated with each option (Fig. 4A). However, whereas a WSLS agent matched perfectly, some IMPs had a slight tendency towards maximizing (Fig. 4A, inset). In order to compare IMPs against a true value-integrating agent, we also simulated matching law behavior from an optimized SARSA agent (*α* = 0.8, *γ* = 0.9). SARSA had a substantially stronger maximizing effect. Thus, although IMPs did have a slight tendency to maximize, they did not maximize to the full extent possible in this task.

**Fig. 4.**
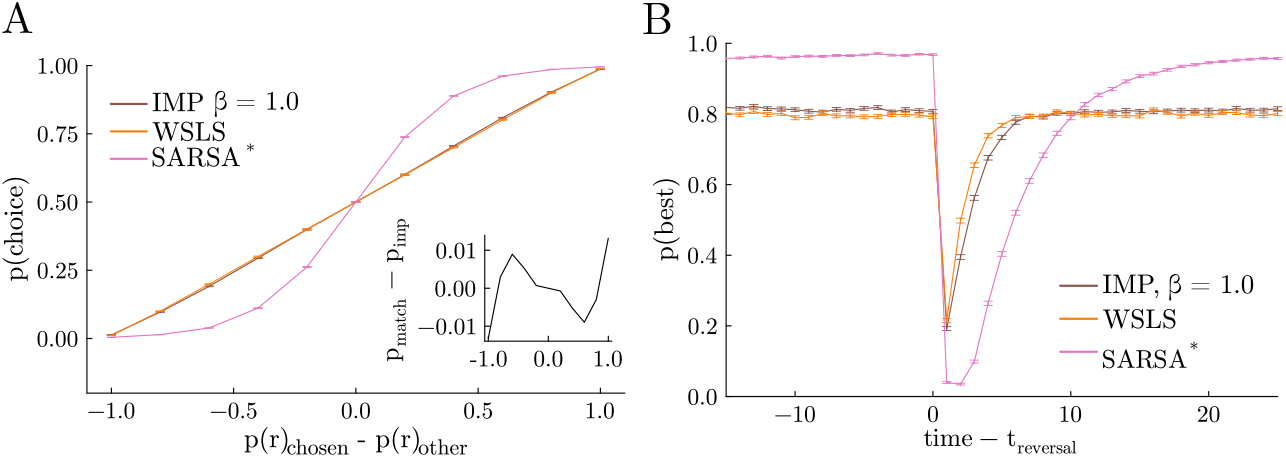
IMPs tend to be matching agents with a slight tendency to maximize reward. (A) Probability of choosing a particular arm as a function of that arm’s probability of reward minus the probability of reward of the unchosen arm. Inset: probability of choice of a perfect matching strategy (a straight line) minus the probability of choice for the IMP agent as a function of the arm’s probability of reward minus the probability of reward of the unchosen arm. (B) Probability of choosing the best arm as a function of time relative to the onset of a reversal event. Tasks described in Sec. 3, with a hazard rate of 0.02. All agents received the same random walks in each session.

### 4.4 IMPs trade-off flexibility and stability

In biological brains, there is a natural trade-off between the ability to persist in stable environments and the ability to adapt to change [33, 5]. In order to determine if IMPs resolve this trade-off, we simulated their behavior in a probabilistic reversal learning task in which stable periods (where one option is clearly more valuable than the other) are interspersed with “reversals” (where the values flip). During stable periods, some IMPs were able to outperform WSLS agents (Fig. 4B) because they were better able to persist in choosing the high-value option despite the noisy reward. By contrast, at reversals, these same IMPs learned less quickly than the WSLS agents. To estimate what a value-integrating agent would do, we again simulated behavior from the optimal SARSA agent (*α* = 0.8, *γ* = 0.4). Although the IMPs were less capable than SARSA during the stable periods state, IMPs adapted faster at reversals—already beginning to reverse after the first omitted reward, whereas SARSA required at least 2 omitted rewards. In sum, IMPs were again able to solve a classic sequential decision-making task and their performance levels were in between the extremes of a memory-less agent and a full reinforcement learning agent.

## 5 Discussion

The ability to robustly store and retrieve information about past interactions with the world is crucial for decision-making. In this paper, we have explored a simple, yet surprisingly competent model for decision-making that incorporates a “faulty” memory. We showed that Imperfect Memory Programs (IMPs) have a memory system that provably trades off the cost of retrieval errors and the cost of high complexity. We found that IMPs do well in classic decision-making tasks under uncertainty, performing similarly or slightly better than agents with a similar strategy but perfect memory. At the same time, IMPs showed characteristics that are more aligned to certain natural and normative decision-makers, such as an exponential reward history kernel, an increased persistence with chosen targets, and a slight tendency to maximize (versus match) rewards.

A memory retrieval system that trades off error and complexity costs also improves performance in structure learning [2]. In that context, imperfect memory can enhance generalization in hierarchically organized environments, task spaces, and other networks because it permits smoothing in learned associations in time. Here, we have applied a similar idea to the notion of reward learning and found that the same errors in retrieval help agents perform a rudimentary form of value integration. Given that human behavior is consistent with hierarchical reinforcement learning [34, 35], imperfect memory may be a good candidate mechanism for smoothing reward associations in such hierarchically organized spaces.

This paper complements recent work on resource-limited decision-making. A particularly relevant line of work is solving a *reward-complexity* trade-off, where a highly complex policy can achieve high rewards, but is cognitively costly [36, 37]. Although the policies derived also involve the Boltzmann distribution, our work aims specifically to solve a error-complexity trade-off in *memory recall*. Combining a resource-limited decision-making system and a resource-limited memory could be an opportunity to offer a parsimonious account of resourcelimited cognitive systems more broadly [38, 39].

Memory limitations have also been explored in the field of reinforcement learning (RL). Because the deep neural networks used in RL are sample inefficient, several groups have begun using episodic memory-based systems to improve learning in data-limited settings [40, 41]. One approach leverages an external memory system to retrieve experiences of high relevance to the agent, in order to update the agent’s policy and value function [42, 43]. In contrast, our approach does not start with the assumption that value is calculated at all. Instead, IMPs demonstrate that samples from episodic memory can be sufficient to imply that value integration is occurring even when it is not.

Although IMPs are simplistic, additional work here could have implications for artificial intelligence research. First, the stochastic retrieval of episodic memories could be extended to actions and states and not just rewards. Agents with stochastic retrieval in all 3 domains would presumably generalize better [2]. Second, retrieving multiple episodic memories might make IMPs even more capable, especially if samples are weighted according to time-relevance or state/action similarity. Overall, given that memory errors are consistent with realistic computations, showing how they can enhance generalization and performance is a promising avenue of research.

## Acknowledgments

Support was provided by the Natural Sciences and Engineering Research Council of Canada (RGPIN-2020-05577), the Research Corporation for Science Advancement & Frederick Gardner Cottrell Foundation (Project 29087), the Canada Research Chair Dynamics of Cognition (FD507106) and a Research Fellowship from the Jacobs Foundation.

## Notes

### Competing Interest Statement

The authors have declared no competing interest.

